# Language learning in a stressful world: individual differences in cardiac responses to stress and recovery predict statistical learning

**DOI:** 10.64898/2026.05.12.724548

**Authors:** Aliva Sholihat, Risto Halonen, Riikka Möttönen, Anu-Katriina Pesonen

**Affiliations:** Cognitive Science, Department of Digital Humanities, Faculty of Humanities, University of Helsinki, Finland; SleepWell Research Program, Faculty of Medicine, University of Helsinki, Finland

**Keywords:** bodily regulation, heart rate variability, individual differences, stress and learning

## Abstract

Learning in adulthood is embedded in everyday social life, in which periods of psychosocial stress alternate with recovery. In such a dynamic environment, the autonomic nervous system regulates bodily responses, including cardiac activity. Individuals differ in their capacity for bodily regulation, but it remains unknown whether these differences modulate statistical learning. Here, we investigated learning of probabilistic patterns in speech streams, i.e., statistical language learning, in a six-hour experiment incorporating psychosocial stress and recovery to approximate everyday conditions. Sixty-five adults were exposed to speech streams in conditions with and without acute stress induction, with learning assessed immediately after exposure and following a rest period. Heart rate variability was recorded throughout the experiment to capture individual differences in cardiac responses across stress and recovery periods. From these measures, we constructed composite proxies of cardiac mobilization during acute stress and cardiac vagal recovery during rest. Individuals with congruent cardiac mobilization and cardiac vagal recovery composite scores, either jointly high or jointly low, showed superior statistical learning outcomes across learning conditions. Cardiac mobilization was positively correlated with immediate performance, i.e., encoding of probabilistic patterns. In contrast, cardiac vagal recovery was positively correlated with post-rest performance, i.e., consolidation. Moreover, the association between momentary cardiac mobilization during speech exposure and statistical learning depended on individuals’ cardiac mobilization composite scores. These findings demonstrate that individual differences in cardiac regulation are closely linked to the ability to learn statistical dependencies in the environment involving changing physiological demands. Overall, the findings highlight the role of bodily regulation in statistical language learning.

## Introduction

Adults frequently experience stress arising from socially evaluative situations, and many learning contexts require acquiring new knowledge under such psychosocial stress ^1,2^. One salient example of such learning context is foreign language learning. For many adult language learners, being immersed in a foreign linguistic and social environment is stressful. The fear of being judged for mistakes or sounding incompetent often evokes self-consciousness and anxiety. This experience, commonly termed as Foreign Language Anxiety ^3^, captures the social-evaluative pressure inherent in language use and, at higher levels, is consistently associated with slower acquisition and poorer learning performance ^4–6^.

Stress is not, however, uniformly detrimental for learning. According to the Yerkes–Dodson principle, performance follows an inverted-U relation with arousal, such that an optimal level of arousal supports learning ^7^. Moderate arousal supports sustaining attention, encoding new information, and promoting neural plasticity ^8,9^. The effects of stress on learning are complex and depend on multiple contextual factors such as its intensity, duration, timing, and the memory systems engaged ^10,11^. While prolonged chronic stress reliably impairs learning ^12^, acute, short-lived stressors show mixed outcomes: in some cases impairing learning, but in others enhancing it ^13,14^.

One potential source of this variability in effects of acute stress on learning lies in individual differences in physiological stress regulation ^10,15^. The autonomic nervous system (ANS), comprising two main interacting branches, the sympathetic nervous system (SNS) supporting mobilization, and the parasympathetic nervous system (PNS) supporting restoration, plays a major role in the rapid physiological response to acute stress ^16,17^. One important target of this regulation is the heart. Autonomic cardiac control refers to the processes through which the sympathetic and parasympathetic influences adjust cardiac activity in response to internal bodily states and changing environmental demands ^18–23^. These influences are expressed, in part, through changes in heart rate and beat-to-beat cardiac timing, which can be examined non-invasively using measures such as heart rate variability (HRV) ^22^.

HRV, which reflects important aspect of autonomic cardiac control, is also closely linked to cognitive functions. The Neurovisceral Integration Model argues for the interplay between autonomic control of the heart with cognitive and emotional regulation involving the interaction of the prefrontal cortex and the central autonomic network processing ^24,25^. Consistent with this framework, studies have shown that individual differences in HRV are associated with executive function, attention, and working memory ^26–28^. Higher resting vagally mediated HRV is associated with better prefrontal regulation, while lower HRV is associated with reduced cognitive flexibility ^26^. Together, this line of evidence suggests that physiological regulation in response to acute stress and the cognitive processes supporting learning may not be independent systems. Thus, these findings motivate investigating whether individual differences in cardiac responses across stress and recovery are associated with variability in learning.

Prior studies have examined how HRV is associated with learning, including in the context of acute stress exposure ^29,30^. However, the importance of HRV to learning in its various forms is not yet well understood ^26,27^. For example, much of the literature has focused on explicit or declarative forms of learning ^14,26^. Explicit learning paradigms rely heavily on conscious strategies, working memory, and executive control — functions that are sensitive to acute stress ^31^. Thus, it is difficult to disentangle the effects of stress on core learning mechanisms from the confounding influence of higher-order cognitive functions. Furthermore, evidence from multiple memory systems framework indicates that acute stress differentially affects learning, often impairing declarative memory while preserving or even biasing behavior toward implicit learning ^32,33^.

Statistical learning—the ability to implicitly extract probabilistic regularities from structured input through passive exposure—is widely regarded as a robust, domain-general and foundational mechanism underlying perception, memory, and learning ^34–36^. Because it does not require explicit instruction or deliberate strategy use, statistical learning provides a powerful framework for examining implicit learning processes. Statistical learning mechanisms play a critical role also in language learning from infancy to adulthood. Statistical language learning (SLL), the ability to detect and memorize hidden reoccurring probabilistic patterns and regularities from linguistic input (e.g., speech streams) based on transitional-probability cues, is widely regarded as a foundational mechanism of language learning ^34,39^. Emerging evidence suggests that while acute stress impairs declarative learning, it may have no effect on or even enhance visual and motor statistical learning, indicating distinct underlying pathways ^37,38^. However, the effects of acute stress on auditory statistical learning and SLL remain largely unknown. Furthermore, the extent to which individual differences in autonomic cardiac responses during stress relate to variability in statistical learning is underdetermined.

In the present study, we aimed to determine how individual differences in cardiac responses to changes in psychosocial environment affect learning the probabilistic regularities from continuous speech streams. More specifically, we asked whether patterns of cardiac responses across repeated stress inductions and subsequent recovery periods can explain variability in (i) SLL outcomes immediately after language exposure and (ii) retention of newly acquired knowledge across a post-learning interval. Rather than assuming that psychosocial stress exerts uniform effects on learning, our approach tested whether a given stressful context may support or hinder learning depending on how individuals regulate physiological mobilization and recovery.

Consistent with the Vagal Tank Theory’s three-Rs ^40^, we considered cardiac responses across three distinct levels: resting, reactivity, and recovery. Resting refers to the cardiac activity during the resting baseline. Reactivity refers to cardiac responses during acute stress events, and recovery refers to post-stress rebound and subsequent opportunities for relaxation and rest. Within our study, we term the stress reactivity dimension as cardiac mobilization (CM), reflecting the expected mobilization of cardiac function to meet environmental demands which typically are expressed through faster heart rate and more constrained beat-to-beat variability ^18,21,22,41^. Whereas for the recovery dimension, we term it as cardiac vagal recovery (CR), reflecting cardiac responses during rest opportunities which typically is expressed through a slower heart rate and greater vagally mediated HRV ^22,42^. We examined individual differences in cardiac response across these dynamics of mobilization of recovery, rather than focusing on physiological stress reactivity at a single time point.

To address our research questions, we used a validated virtual-reality adaptation of the Trier Social Stress Test (VR-TSST) ^43^, supplemented with Virtual Reality Stroop Task ^44^, which altogether induced repeated psychosocial stress in participants. We extended the standard TSST protocol with guided relaxation (Yoga Nidra) followed by a 90-minute nap or wakeful rest, to approximate the dynamics of stress and recovery in everyday life. Participants listened to novel speech streams with hidden probabilistic patterns (i.e., artificial languages) under with-stress-induction and no-stress-induction learning contexts. They completed a two-alternative forced-choice (2AFC) task immediately after exposure and after rest periods, allowing us to assess acquisition and retention of new linguistic knowledge. To measure cardiac responses across the experiment, we continuously recorded heart-rate variability (HRV) and supplemented this with electrodermal activity (EDA) as a more direct measure of the stress response ^21,25^. By integrating naturalistic psychosocial stress induction with continuous cardiac recording and a well-established auditory SLL paradigm, we were able to examine how bodily regulation in a stressful environment shapes the ability to acquire new knowledge from speech streams.

## Results

### Verification of psychosocial stress induction

Before analyzing statistical learning outcomes, we verified that the stress inductions elicited the expected physiological stress responses. In a within-subject, counterbalanced design, participants completed statistical language learning (SLL) tasks under both with-stress-induction and no-stress-induction learning contexts (Fig. 1A and 1B). Participants were assigned to one of two counterbalanced order groups. The AB group completed the statistical language learning (SLL) task first under the no-stress-induction context and later under the with-stress-induction context, whereas the BC group completed the tasks in the reverse order.

**Fig. 1.**
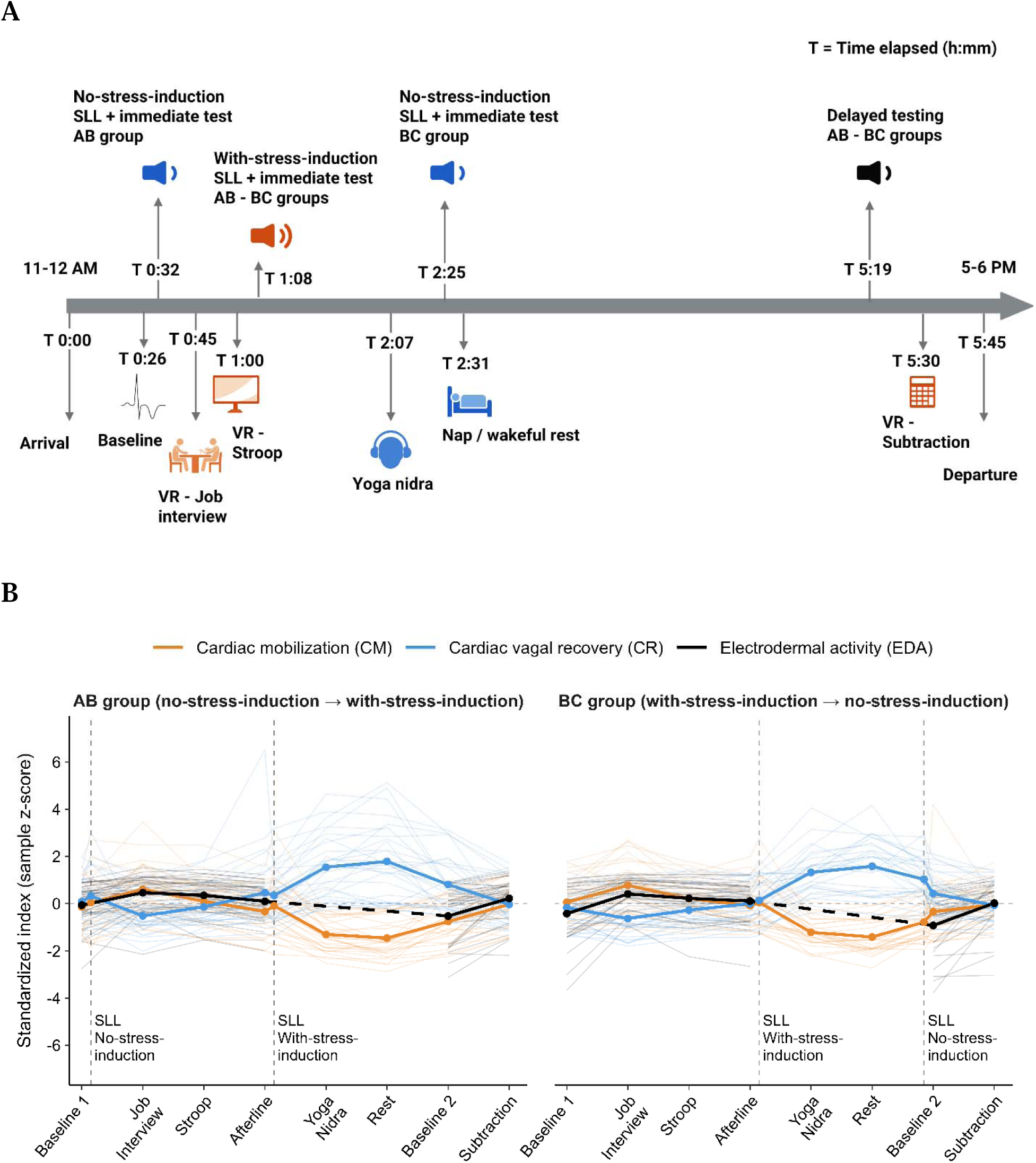
(A). Overview of experimental timeline. Participants completed statistical language learning (SLL) under no-stress-induction and with-stress-induction contexts in one of two counterbalanced orders (AB or BC). The AB group completed SLL task first under the no-stress-induction context and later under the with-stress-induction context, while the BC group completed the tasks in the reverse order. The protocol incorporated a modified Virtual Reality Trier Social Stress Test (VR-TSST), including an additional VR Stroop task, and an extended recovery period comprising 15 min of Yoga Nidra followed by a 90-min nap or wakeful rest. Only the main experimental events are shown, additional procedures are omitted for clarity (e.g., post-task measurements, subjective affect questionnaires, snack time, and shower time). Timestamps represent approximate elapsed times from arrival. Participants spent approximately six hours in the experiment. **(B). Physiological measures dynamics during the six-hour experiment.** Cardiac mobilization (CM; orange), cardiac vagal recovery (CR; blue), and electrodermal activity (EDA; black) indices are plotted across experimental tasks. While the HRV-derived cardiac measures were recorded continuously, EDA was recorded during the baseline and stress-induction tasks but not during SLL, Yoga Nidra, or the rest period. For visual comparison, all three physiological measures were z-standardized within the study sample. The group-level trajectories confirmed effective stress induction, with both CM and EDA peaking during the Job Interview task. Effective recovery was reflected in reduced CM and increased CR, with the latter peaking during the rest period of nap/wakeful rest. Thin lines represent individual trajectories, whereas thick lines represent group means. Marked interindividual variability in response magnitude underscores the importance of modelling physiological dynamics at the individual level rather than relying solely on group averages.

Natural-log-transformed skin conductance level measured via electrodermal activity increased significantly during the stress induction tasks relative to baseline: Job Interview, *t*(59) = 7.77, *P* < .001, Cohen’s *d*_z_ = 1.00; Stroop, *t*(58) = 7.13, *P* < .001, Cohen’s *d*_z_ = 0.93; Subtraction, *t*(59) = 8.69, *P* < .001, Cohen’s *d*_z_ = 1.12. Mean heart-rate change from baseline during learning was also significantly higher in the with-stress-induction learning context than in the no-stress-induction learning context, *t*(59) = 4.55, *P* < .001, Cohen’s *d_z_* = 0.59.

When the counterbalanced task-order groups (AB and BC) were combined, cardiac mobilization (CM) index was higher in the with-stress-induction than no-stress-induction learning context, *t*(59) = 3.04, *P* = .004, Cohen’s *d*_z_ = 0.39. However, the magnitude of this difference depended on task order. In the BC group, average CM index was greater during with-stress-induction than during no-stress-induction learning (*t*(28) = 9.03, *P* < 0.001, Cohen’s *d*_z_ = 1.68). Meanwhile, in the AB group, average CM index during no-stress-induction learning was comparable to that observed during subsequent with-stress-induction learning (*t*(30) = -1.92, *P* = 0.065, Cohen’s *d*_z_ = -0.34), consistent with elevated anticipatory arousal at experimental onset. Together, these convergent measures confirm physiological activation during stress-induction tasks and greater cardiac mobilization during with-stress-induction learning overall, while indicating that physiological differentiation between the learning contexts depended on task order (AB or BC).

### No differences in SLL between with-stress-induction and no-stress-induction learning contexts

We next examined group-level SLL outcomes (Fig. 2). Participants were exposed to a two-minute artificial language stream under with-stress-induction and no-stress-induction learning contexts and completed two-alternative forced-choice (2AFC) tests immediately after exposure and again after rest periods (either wakeful rest or daytime nap).

**Fig. 2.**
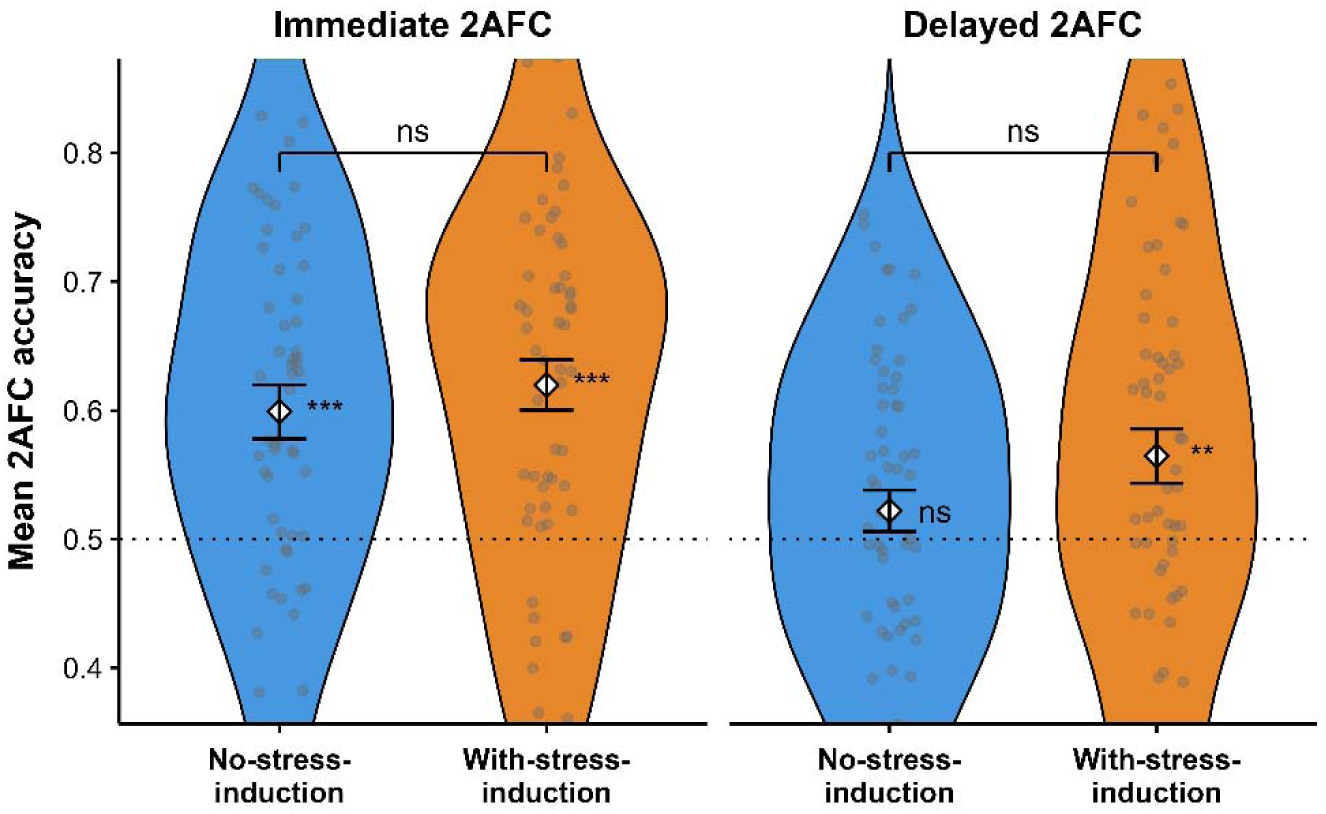
Mean two-alternative forced-choice (2AFC) accuracy in immediate and delayed memory tests for language exposures under no-stress-induction and with-stress-induction learning contexts. Participants performed significantly above chance in the immediate memory tests, indicating learning in both no-stress-induction and with-stress-induction learning contexts. In the delayed memory tests, participants performed above chance only for the language which they had been exposed to under with-stress-induction context. Overall, different stress contexts during language exposure did not significantly affect accuracy at either phase. Violin plots display individual data points, group means (white diamonds), and error bars represent ±1 SEM across participants (N = 60).

Accuracy exceeded chance (.50) in the immediate test in both with-stress-induction (*M* = .62, *t*(59) = 6.14, *P* < .001) and no-stress-induction learning contexts (*M* = .60, *t*(59) = 4.74, *P* < .001), indicating successful initial learning (Fig. 2). In the delayed memory test, accuracy remained above chance for language exposed in the with-stress-induction learning context (*M* = .56, *t*(59) = 3.05, *P* = .003) but not for language exposed in the no-stress-induction learning context (*M* = .52, *t*(59) = 1.35, *P* = .18). Direct comparisons between with-stress-induction and no-stress-induction learning contexts, however, revealed no significant differences at either the immediate test (*t*(59) = −0.67, *P* = .50) or delayed test (*t*(59) = −1.51, *P* = .14).

These results suggest that participants acquired new linguistic knowledge equally effectively in both no-stress-induction and with-stress-induction learning contexts. Since there was pronounced interindividual variability in cardiac responses to stress and recovery (Fig. 1B), further analyses focused on the role of individual differences in cardiac response for the learning outcomes.

### Construction of individual-level cardiac response composites

To capture individual characteristics of cardiac response patterns across the six-hour experiment, we constructed separate composite scores for cardiac mobilization (CM) and cardiac vagal recovery (CR). We first converted segment-level CM and CR indices into baseline-referenced change/delta scores. We then applied separate principal component analysis (PCA) to theoretically group delta scores from stress-induction and recovery periods.

The CM composite was based on delta scores during the stress induction tasks of Job Interview, Stroop Test, and Subtraction Task. Its first principal component explained an average of 59.1% of variance (95% CI: 51.8–66.2%). The CR composite was based on delta scores during Yoga Nidra, Nap or Wakeful Rest, and Post-Job Interview Rebound and explained 62.7% of variance (95% CI: 55.1–73.3%). All tasks contributed meaningfully, with the strongest weights observed for Job Interview (40.6%) and Yoga Nidra (41.3%).

The individual CM composite and CR composite scores were later used as participant-level predictors in the mixed-effects model. These participant-level composites were distinguished from learning-period CM, which refers to the CM index measured during each speech-stream exposure (with-stress-induction learning and no-stress-induction learning) and was entered as a context-specific predictor in the model.

### Congruency of cardiac mobilization and cardiac vagal recovery supports statistical learning

A generalized linear mixed-effects model (GLMM) predicting trial-level accuracy (*N* = 3,840 trials; 60 participants) in the SLL task revealed several significant effects (Supplementary Table S1). Accuracy was lower at delayed memory test (OR = 0.74, 95% CI [0.65–0.85], *P* < 0.001), consistent with partial forgetting over time. Acute stress induction did not significantly influence the probability of a correct response (OR = 1.15, 95% CI [0.93–1.43], *P* = 0.206), in line with the group level analyses result (Fig. 2).

The CM and CR main effect terms were not statistically significant (CM: OR = 1.03, 95% CI [0.95–1.13], *P* = 0.468; CR: OR = 1.07, 95% CI [0.98–1.16], *P* = 0.154). In contrast, the interaction between CM and CR composites was significantly associated with learning accuracy (OR = 1.15, 95% CI [1.05–1.25], *P* = 0.002). The probability of a correct response was higher when CM and CR composites were congruent (i.e., aligned in the same direction). This is illustrated in (Fig. 3A) showing that congruent profiles, i.e., jointly high or jointly low CM-CR composite profiles were associated with higher learning accuracy. This congruence effect was evident across both immediate and delayed memory tests as testing phase did not significantly moderate the effect (*P* = 0.21).

**Fig. 3.**
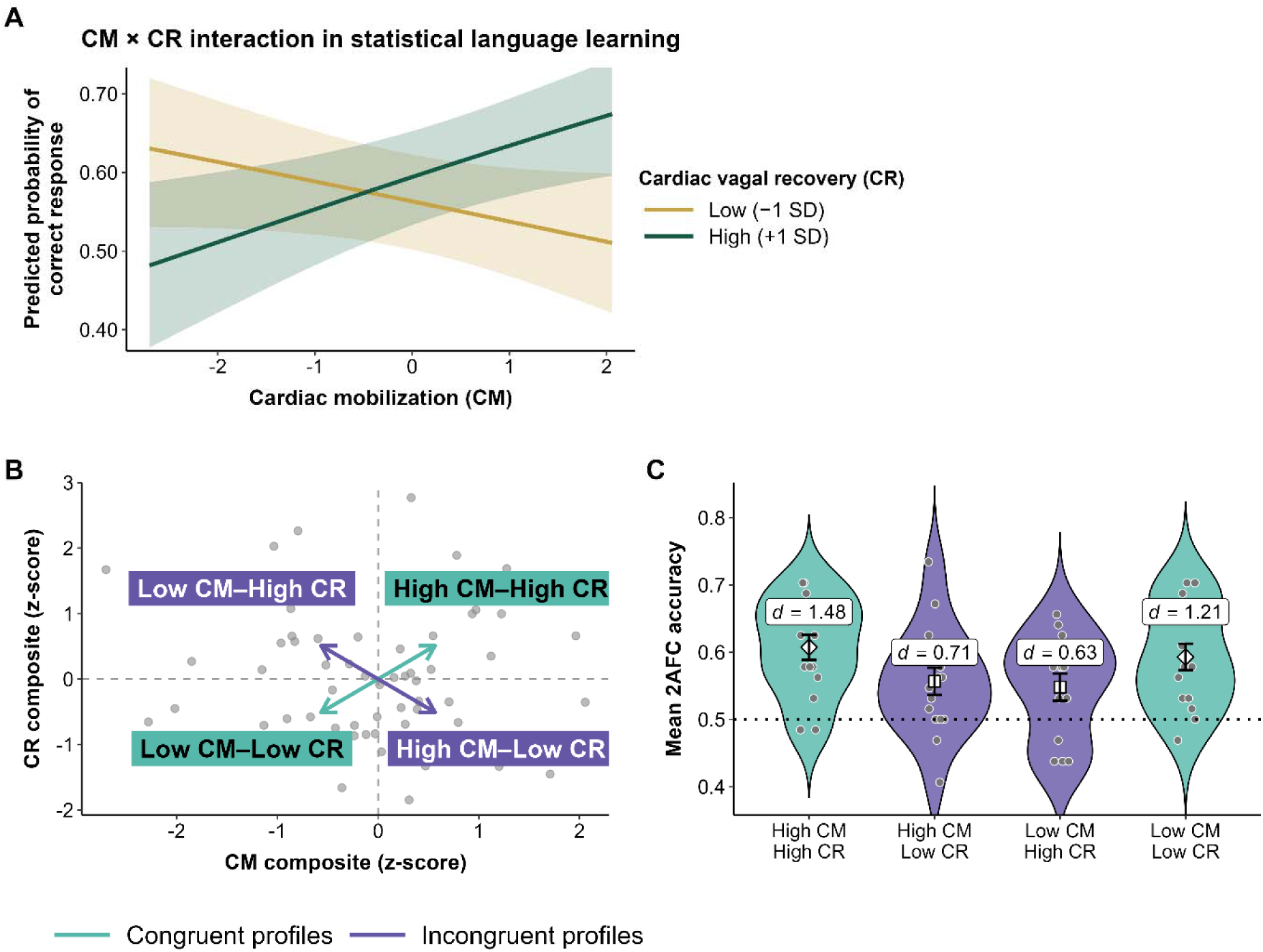
Cardiac response profiles predict statistical language learning (SLL) accuracy. **(A).** Model-predicted interaction between cardiac mobilization (CM) and cardiac vagal recovery (CR) composites in statistical language learning. CM and CR composites summarize each participant’s typical pattern of cardiac response across the tasks in this experiment: how strongly their heart responded to stress inductions (CM) and to opportunities for rest and recovery (CR), relative to baseline. The predicted probability of a correct response is shown as a function of CM composite at low (−1 SD) and high (+1 SD) levels of CR composite, illustrating cardiac response congruence. Predicted accuracy is higher when CM and CR composite scores are congruent (jointly high or jointly low) than when they are incongruent. Shaded areas indicate 95% confidence intervals. **(B).** Scatterplot of individual participants within the cardiac response space defined by z-scored cardiac mobilization (CM) and cardiac vagal recovery (CR) composite scores. Each grey dot represents one participant positioned according to their CM and CR composite scores. Profiles were defined via mean splits (0 = sample mean), providing a categorical visualization of the continuous CM × CR interaction observed in the mixed-effects model. Two congruent profiles (High CM–High CR; Low CM–Low CR) and two incongruent profiles (High CM–Low CR; Low CM–High CR) are shown. Arrows illustrate congruent versus incongruent cardiac response patterns across CM and CR axes. **(C).** Mean two-alternative forced-choice (2AFC) accuracy across cardiac response profiles, averaged across memory testing phases and stress contexts. Descriptively illustrating the modeled interaction, congruent profiles (High CM–High CR; Low CM–Low CR) outperformed incongruent profiles in SLL. Violin plots show participant distributions; grey points indicate individual means; black points and bars denote group means ± SEM. Dashed line indicates chance level accuracy (0.5). Cohen’s d values indicate effect sizes relative to chance level accuracy.

To facilitate interpretability, we visualized this continuous model-based interaction effect descriptively using categorical cardiac response profiles. Participants were classified via mean splits of CM and CR composites (z-scored; 0 = sample mean), yielding two congruent (High CM–High CR; Low CM–Low CR) and two incongruent cardiac response profiles (High CM–Low CR; Low CM–High CR) (Fig. 3B).

All profiles performed above chance, yet mean accuracy varied systematically (Fig. 3C). Congruent cardiac response profiles showed the highest accuracy (High CM–High CR: *M* = 0.61; Low CM–Low CR: *M* = 0.59), whereas incongruent cardiac response profiles showed lower accuracy (High CM–Low CR: *M* = 0.56; Low CM–High CR: *M* = 0.55). Thus, the categorical visualization converged with the continuous GLMM findings: individuals exhibiting congruent CM–CR profiles demonstrated superior SLL (*P* = 0.01, *d* = 0.63).

### Cardiac mobilization preferentially supports encoding and cardiac vagal recovery preferentially supports consolidation

Learning was assessed in two phases: immediately after exposure, as a proxy to the encoding phase, and approximately three hours later after a rest period (nap or wakeful rest), as a proxy to the consolidation phase. The GLMM revealed significant phase-dependent associations between the cardiac response and probability of correct response (i.e., language learning). A significant CM composite × Phase interaction (OR = 0.83, 95% CI [0.72–0.95], *P* = 0.008) indicated that higher CM composite score was associated with a higher accuracy in the immediate memory test, whereas a significant CR composite × Phase interaction (OR = 1.17, 95% CI [1.02–1.34], *P* = 0.025) indicated that higher CR composite score was associated with higher accuracy in the delayed memory test (Fig. 4A and 4B).

**Fig. 4.**
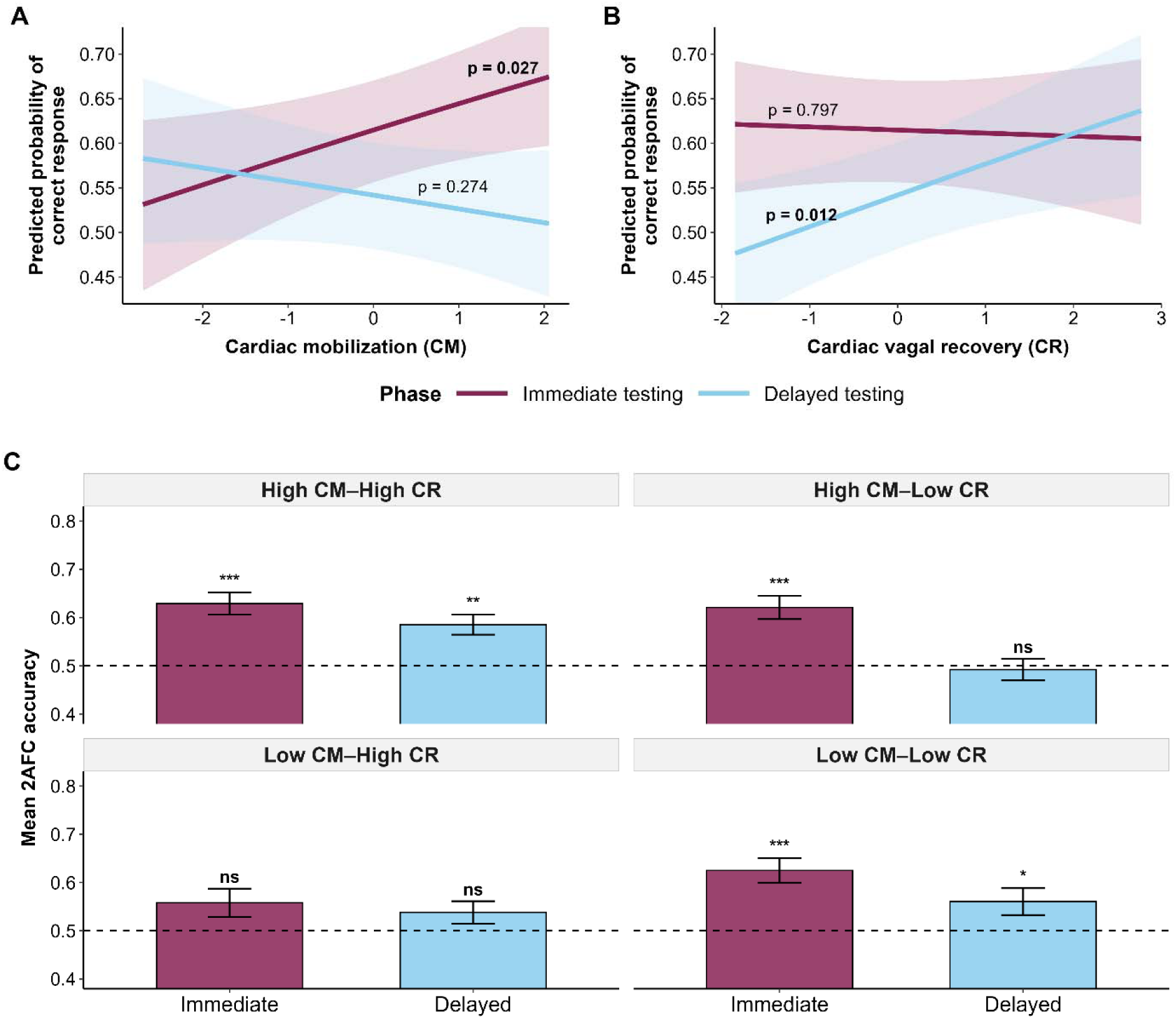
Phase-dependent associations of cardiac mobilization (CM) and cardiac vagal recovery (CR) composite scores with statistical learning. **(A).** Model-predicted probability of a correct response as a function of CM composite during immediate and delayed testing. Higher CM composite scores were associated with greater immediate-test accuracy (slope *P* = 0.027). **(B).** Model-predicted probability of a correct response as a function of CR composite during immediate and delayed testing. Higher CR composite scores were associated with greater delayed-test accuracy (slope *P* = 0.012). Burgundy and light-blue lines represent immediate and delayed testing, respectively. Shaded regions indicate 95% confidence intervals. Displayed *P* values correspond to phase-specific simple-slope tests. **(C).** Phase-specific accuracy across cardiac response profiles. Mean two-alternative forced-choice (2AFC) accuracy during immediate (encoding) and delayed (post-rest consolidation) testing across cardiac response profiles. The incongruent High CM–Low CR profile showed significantly above-chance accuracy at the immediate test but did not differ from chance at delayed testing. The incongruent Low CM–High CR profile did not differ from chance at either phase. In contrast, congruent profiles (High CM–High CR; Low CM–Low CR) maintained stable, above-chance accuracy across testing intervals. Bars represent group means ± SEM. The dashed horizontal line indicates chance level accuracy (0.50). Significance markers denote comparisons against chance within each phase (\**P* < 0.05, ** *P* < 0.01, *** *P* < 0.001; ns, not significant).

To characterize these continuous phase-dependent coordination effects descriptively, we examined mean 2AFC accuracy across phases in different cardiac response profiles (Fig. 4C). In incongruent High CM–Low CR profile, mean accuracy was above chance level in the immediate 2AFC test, but did not differ from chance level in delayed 2AFC test, consistent with the idea that high CM composite supports encoding, while low CR composite impairs consolidation. In the incongruent Low CM–High CR profile, mean accuracy did not differ from the chance level at either phase, consistent with the idea that low CM composite impairs encoding. Notably, participants in the congruent cardiac response profiles maintained stable, above-chance accuracy across immediate and delayed 2AFC tests.

### The effect of momentary cardiac mobilization on statistical learning depends on cardiac mobilization response profile

Next, we examined how CM composite profiles interacted with momentary CM during language exposure. The GLMM (Supplementary Table S1) revealed a significant interaction between CM composite and momentary CM during language exposure (OR = 1.16, 95% CI [1.06–1.27], *P* < 0.001). This indicates that the effect of momentary CM during language exposure was not uniform across individuals, but varied as a function of their characteristic CM composite profiles. Model-predicted probabilities (Fig. 5) showed that, among participants with lower CM composite, higher momentary CM during language exposure was associated with lower predicted accuracy. In contrast, among participants with higher CM composite, higher momentary CM was associated with higher predicted accuracy.

**Fig. 5.**
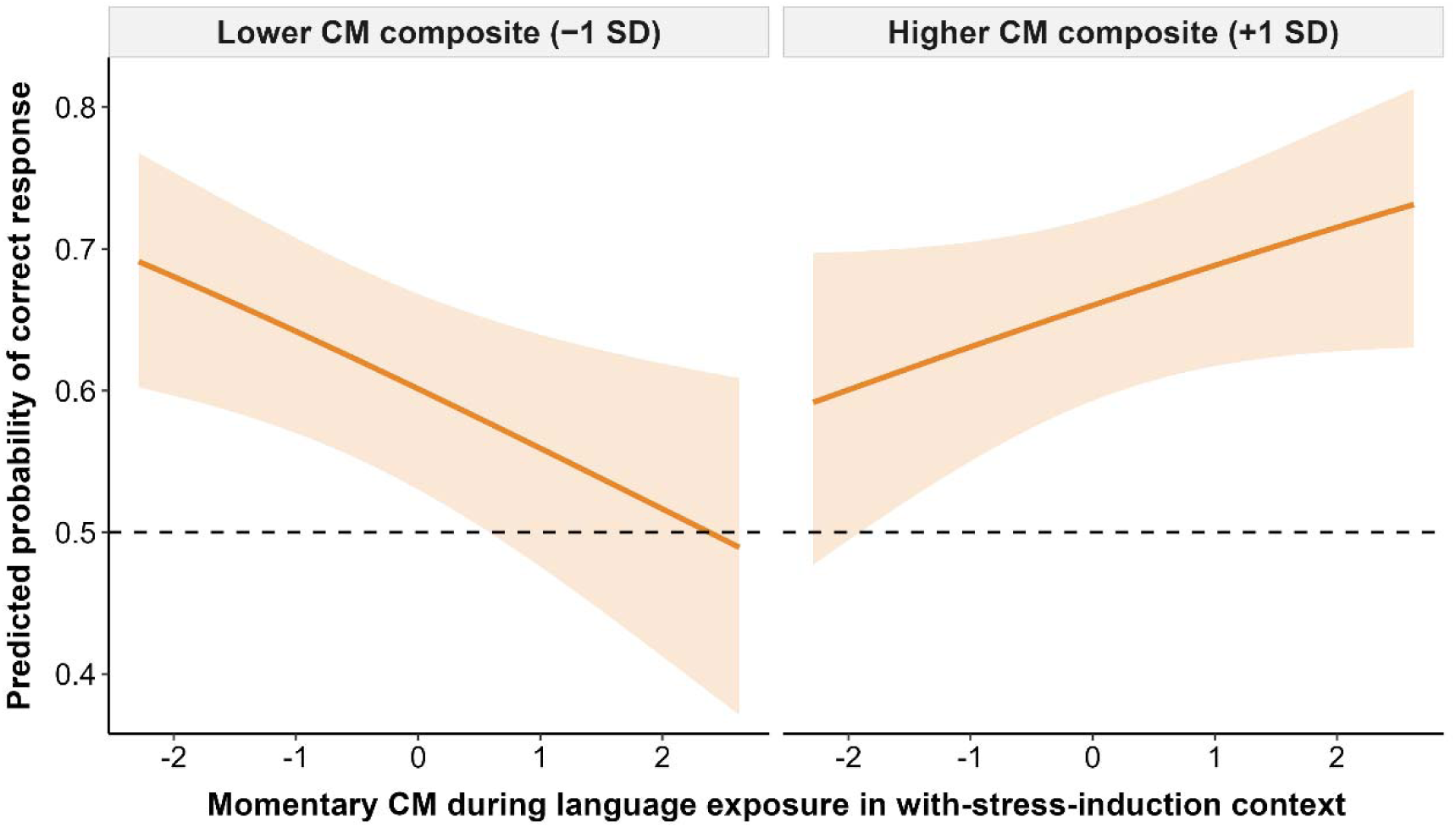
Model-predicted association between momentary cardiac mobilization (CM) during language exposure and learning accuracy as a function of CM composite profile. Predicted probability of a correct response from the GLMM is shown for the with-stress-induction learning context, separately for participants with lower and higher CM composite profiles. Shaded areas indicate 95% confidence intervals. The dashed line indicates chance-level performance. The direction of the association between momentary CM and predicted accuracy differed by CM composite profile.

### Post-hoc sensitivity analyses

To address concerns about the transparency of Kubios’s proprietary scoring, we fitted linear regression models using the publicly documented R–R-derived components of its indices. These models closely reproduced Kubios outputs (SNS: R² = .9998, RMSE = .0190; PNS: R² = .9999, RMSE = .0107), providing transparent approximations from which CM and CR indices could be constructed using accessible R–R-derived measures (Supplementary Fig. S1).

We also tested alternative operationalizations of CM and CR using different combinations of conventional cardiac measures. Key GLMM interactions remained significant across several tested alternatives, suggesting that these findings were not solely dependent on Kubios’s proprietary scoring, although statistical support varied across operationalizations (Supplementary Table S2). However, key CM and CR interaction was not significant when both constructs were operationalized as RMSSD withdrawal and RMSSD increase, respectively (*P* = 0.104, Supplementary Table S2).

Furthermore, we added baseline measures as covariates in the final model to test whether the associations remained after accounting for individual differences in initial resting cardiac measures (Supplementary Table S3). Our original operationalization with Kubios-derived CM and CR retained all four significant interactions after controlling for baseline (Supplementary Table S4). Among the alternative operationalizations, using mean HR as a proxy for CM and an equally weighted combination of standardized mean RR interval and raw RMSSD as a proxy for CR also retained all four significant interactions (Supplementary Table S5-6). Full methods and results of post-hoc analyses are provided in the Supplementary Material.

## Discussion

Here, we investigated how individual differences in autonomic cardiac responses were associated with SLL in a dynamic environment with varying acute stress and recovery periods approximating real-life conditions. Using a six-hour experimental paradigm that included repeated naturalistic psychosocial stress inductions, recovery periods, and an auditory SLL task, we examined how bodily regulation affects the acquisition of knowledge from speech streams with hidden probabilistic structures. We observed significant variation across individuals in their cardiac mobilization (CM) during stress and cardiac vagal recovery (CR) during rest periods. These individual differences in coordination of mobilization and recovery were associated with statistical learning outcomes across learning in no-stress-induction and with-stress-induction context, as well as how momentary cardiac mobilization during speech exposure modulated learning in the with-stress-induction context.

Prior work has often emphasized cardiac vagal recovery, typically indexed through HRV–derived vagal measures, as markers of regulatory flexibility and adaptive capacity ^24,26,28^. Our findings are broadly consistent with HRV-based accounts in showing that autonomic cardiac control plays an important role in cognitive functions ^26,27^, but we also extend this literature by suggesting that successful learning depends not on cardiac vagal recovery alone, but on the coordinated interplay between mobilization and recovery.

To the best of our knowledge, our study provides the first empirical evidence that the interaction between cardiac mobilization during acute stress and subsequent cardiac vagal recovery during rest is associated with auditory statistical language learning. Our finding can be embedded in the Vagal Tank Theory, providing a theoretical basis for examining joint stress reactivity–recovery patterns that may reveal aspects of adaptive regulation not evident when either phase is considered independently ^40^. However, prior empirical studies which directly tested whether stress reactivity and subsequent recovery interact in predicting learning remain scarce ^45–48^.

Our study adds to the literature by highlighting a possible limitation of current operationalizations of stress reactivity and recovery, which commonly use a single vagally mediated HRV metric such as RMSSD (root mean square of successive differences) ^49^. Stress reactivity or cardiac mobilization is often operationalized as a decrease in RMSSD (withdrawal), while recovery is operationalized as an increase in RMSSD ^40,49^. We tested this common approach in our model but did not find evidence of significant interaction. In contrast, with multiparametric representations combining complementary features of the R-R signal, the interaction was evident and the model fit improved. It is plausible that these multiparametric representations provide additional information about cardiac response dynamics during stress and recovery beyond what can be captured by a single vagally mediated HRV metric ^50–53^.

Our two-dimensional model of cardiac mobilization (CM) and cardiac vagal recovery (CR) can be represented by four quadrants, corresponding to low CM–high CR, low CM–low CR, high CM–high CR, high CM-low CR cardiac response profiles. We found that individuals with congruent CM and CR profiles (either both high or both low) outperformed individuals with incongruent profiles in learning probabilistic patterns from speech streams.

Among the incongruent profiles, our findings suggest that their learning bottlenecks may occur at different phases. Participants with high CM but low CR showed relatively preserved immediate learning but a marked decline at delayed memory test, consistent with the possibility that insufficient cardiac vagal recovery at rest compromised post-encoding stabilization. Conversely, participants with low CM but high CR performed poorly at both phases, raising the possibility that weaker initial encoding limited any subsequent benefit associated with greater cardiac vagal recovery.

In contrast, participants with congruent cardiac response profiles maintained stable accuracy across learning phases. This pattern suggests that effective learning may depend on coordinated autonomic cardiac control across distinct stages of the learning process ^24,40,54^. Adequate cardiac mobilization may support attention and encoding during exposure ^55^, while sufficient cardiac vagal recovery may be required to stabilize and retain newly encoded representations ^47^. We interpret these findings as suggesting possible differences in the functional characteristics of the four cardiac response profiles, as summarized in Table 1.

**Table 1.** Proposed functional interpretation of cardiac response profiles and their associations with statistical language learning.

| **Cardiac response profile** | **Proposed functional interpretation** | **Descriptive learning pattern** |
| --- | --- | --- |
| High CM – High CR | Dynamic flexibility | Efficient learning |
| Low CM – Low CR | Stable regulation | Efficient learning |
| High CM– Low CR | Mobilization dominance, insufficient recovery | Less efficient consolidation |
| Low CM – High CR | Recovery dominance, insufficient mobilization | Less efficient encoding |

*Note.* CM = cardiac mobilization; CR = cardiac vagal recovery. Functional labels and associated learning patterns are interpretations of the study’s findings, rather than direct measurement of the mechanisms.

However, it is important to highlight that we do not have sufficient evidence to claim that cardiac response profiles represent stable long-term traits. The CM and CR composites captured coherent individual differences across multiple stress inductions and recovery opportunities, but they were only observed within the six-hour experimental period. It is also possible that these cardiac response profiles represent distinct regulatory modes that individuals dynamically enter depending on changing contexts. For example, individuals may shift between congruent High CM – High CR or Low CM – Low CR according to the level of current environmental demands. Meanwhile, factors such as fatigue, lack of sleep, boredom, or prolonged stress may shift individual’s regulatory mode to incongruent profiles. Another possible interpretation is that better learners are not characterized by a fixed cardiac response profile, but by a greater tendency to enter a congruent regulatory mode (either High CM – High CR or Low CM – Low CR) appropriate to the situation. Longitudinal and repeated-measures studies across different stress and recovery contexts are needed to test these different interpretations.

Our study also contributes empirical evidence on how acute stress affects statistical learning. Previous work has suggested that acute stress may enhance statistical learning by shifting memory systems away from model-based explicit learning toward more model-free implicit learning pathways ^37,38,56,57^. However, our findings demonstrate that the effects of psychosocial stress on statistical learning are not uniform across individuals. We found no differences in statistical learning between the with-stress-induction learning context and the no-stress-induction learning context.

Importantly, we found that elevated cardiac mobilization during language exposure was associated with better learning outcomes in individuals with higher cross-tasks CM composite scores, but with poorer learning outcomes in individuals with lower cross-tasks CM composite scores. This suggests that a similar level of cardiac mobilization (i.e., physiological stress response) can have different functional significance depending on the individual’s characteristic cardiac mobilization profile. This may help explain not only the absence of a group-level effect of stress context in our study, but also why prior studies of acute stress and learning have often reported mixed or inconsistent group-level findings ^14^. Our findings suggest that whether acute psychosocial stress enhances, impairs, or does not affect SLL depends on individual differences in bodily regulation in a stressful environment.

Building on these empirical findings, we draw on allostasis and the predictive processing framework ^58–63^ to speculate why the ability to learn probabilistic patterns from sensory input is associated with individual differences in bodily regulation, especially the congruence of cardiac responses between mobilization and recovery in a dynamic environment. Autonomic cardiac control may be linked to statistical learning through overlapping systems that regulate bodily state, arousal, and sensory processing ^24,25^.

Allostasis refers to the process by which an organism anticipates bodily needs and prepares physiological and behavioral responses to meet them. It describes how the organism achieves stability through adaptation and change ^62,63^. Predictive processing framework offers a conceptual account of how this process may occur. The brain uses learned models to predict future demands or upcoming information, while differences between expected and observed signals generate prediction errors that help update those models ^59^. Learning therefore is important because by remembering outcomes, the brain can use past experiences to improve future anticipation and avoid repeating costly errors. Statistical language learning (SLL) is a specific example of acquiring knowledge from sensory experience: humans extract regularities from speech that can inform expectations about upcoming sensory input ^64^. Thus, SLL can be situated within the broader acquisition of environmental knowledge that supports prediction and adaptation. In particular, when learning occurs under a stressful context where uncertainty is higher and energetic demands increase, this adds further complexity to the prediction the brain needs to process ^61^.

In the brain, these processes of bodily regulation and learning from sensory input may involve overlapping systems. Autonomic cardiac control, through which the sympathetic and parasympathetic influences adjust cardiac activity in response to internal bodily states and changing environmental demands is one possible pathway through which anticipatory regulation is implemented. The prefrontal cortex interacts with the central autonomic network, affecting autonomic control of the heart as well as cognitive and emotional regulation ^24,25^. Neuromodulatory systems involved in autonomic cardiac control provide a further plausible link between bodily regulation and learning. Noradrenergic activity during salient or demanding events modulates attention, sensory processing, and memory, linking physiological mobilization with encoding of new information ^9^. Meanwhile, reduced central cholinergic activity during rest facilitates hippocampal-cortical communication crucial to memory consolidation ^65^.

Allostatic theory also suggests that efficient regulation depends on matching physiological responses to expected demands, together with complementary periods of mobilization and recovery as those demands change ^62,63^. A fixed level of physiological responses would be excessive and wasteful for smaller demands, but insufficient for larger demands ^62,63^. In the context of learning, this suggests that cardiac mobilization (CM) during periods requiring attention and encoding may need to be coordinated with subsequent cardiac vagal recovery (CR), where energetic demands decrease and memory can be consolidated. We propose that our finding of CM-CR congruence across the dynamics of stress and recovery is consistent with this allostatic principle: individuals with congruent cardiac responses across mobilization and recovery showed efficient statistical learning. Nevertheless, this mechanistic interpretation remains speculative and needs to be directly tested in future studies.

In conclusion, our study suggests that individual differences in bodily regulation are tightly linked to the ability to learn patterns and regularities from external sensory signals in information-rich and socially demanding contexts. Statistical learning was optimized when cardiac responses between mobilization during stress and vagal recovery during rest were congruent—either jointly high or jointly low—and when momentary cardiac mobilization during learning aligned with an individual’s underlying cardiac mobilization profile. These findings demonstrate that statistical learning, a core cognitive process, is supported by dynamic interactions between the brain, the body, and the environment.

### Limitation and future directions

Several limitations of the present study should be acknowledged. First, due to practical instrumentation constraints in our extended experiment, we did not measure respiration. RMSSD as the main HRV measure component in our cardiac vagal recovery (CR) index is argued to be less sensitive to respiratory variation, but it remains influenced by breathing rate and depth ^22,49^. This is particularly relevant for our experimental tasks which included speaking, relaxation with Yoga Nidra, and sleep. Thus, changes in breathing rate may have contributed to the cardiac patterns. Using a multiparametric index reduces reliance on a single HRV metric, but it cannot fully eliminate respiratory noise. Future studies using extended naturalistic designs may consider ambulatory instrumentation that can measure both ECG and respiration.

Second, this study focused on the rapid autonomic cardiac control component of the stress response, which is associated with the SAM (sympatho-adreno-medullary) axis rather than the slower HPA (hypothalamic-pituitary-adrenal) axis. Therefore, the effect of cortisol release on the observed effects cannot be determined. We distinguished stress and recovery periods primarily through cardiac indices; however, these measures cannot establish whether cortisol had returned to baseline, and carryover from preceding stress tasks remain possible. For a more comprehensive investigation, future studies can include both axes of stress response.

Next, reliance on Kubios-derived measures with proprietary weighting may limit transparency and reproducibility of our CM and CR indices. We mitigated this issue by conducting a post-hoc linear reconstruction of the Kubios-derived indices from their publicly documented R-R components. While the reconstructed indices showed excellent fit, external validation is still needed to establish the generalizability of our regression model. We also triangulated the results using alternative transparent operationalizations. A model in which mean-HR represented CM and a combination of mean R-R interval with RMSSD represented CR produced consistent findings. These analyses partly addressed the transparency concern and suggest that the findings were not dependent on Kubios-derived indices. However, both the reconstructed and alternative indices were tested post-hoc and require replication in independent studies.

Importantly, due to the limitation of interpretation from R-R interval signals, in this study we examined autonomic cardiac control without making any branch-specific claim of sympathetic or parasympathetic influence. Especially during cardiac mobilization, sympathetic activation and parasympathetic withdrawal cannot be disentangled from R-R derived measures alone ^66^. Future studies could include more physiologically selective measures of cardiac contractility associated with sympathetic beta-adrenergic activity, such as pre-ejection period (PEP) ^67^. However, PEP requires impedance cardiography, which may present additional instrumentation challenges for a naturalistic extended stress-recovery paradigm.

Another limitation concerns the scope of the present experimental paradigm. The findings are specific to acute psychosocial stress and to an implicit SLL task with relatively brief exposure. They may therefore not apply to chronic stress conditions or to learning domains that rely more on explicit or strategic processes. Extending this framework across stress timescales and learning systems will be necessary to determine its generalizability.

Despite these limitations, the concept of cardiac response congruence as the body moves between stress and recovery provides a testable framework for understanding variability in learning under stress. Moderate psychosocial stress is common in educational, occupational, and performance settings, including in the context of language learning, yet its effects on learning remain difficult to predict. Our findings suggest that variability in learning outcomes may partly reflect differences in how individuals physiologically mobilize and regulate arousal across contexts. From this perspective, tailoring learning environments or regulation strategies to individuals’ cardiac response profiles represents a promising direction for future translational research.

## Materials and methods

### Participants

We recruited participants through on-campus advertisements across several universities in the Helsinki region and via online channels. A total of 65 adult Finnish speakers (aged 19–48 years; *M* = 24.6) took part in the study. All participants were generally healthy, the exclusion criteria being presence of sleep disorders, using medication affecting sleep, and hearing problems. Seven participants self-reported a neurodevelopmental or learning disorder (e.g., ADHD or dyslexia), and six were left-handed or ambidextrous. Although SLL studies often exclude individuals with these characteristics, inclusion criteria were permissive due to the complexity of the multiparadigm design and recruitment constraints. The sample comprised 66% female and 34% male participants. After excluding outliers with extreme cardiac mobilization or cardiac vagal recovery responses using IQR (inter-quartile range) filtering, the final dataset included 60 participants. The study was approved by the University of Helsinki Ethics Committee and conducted in accordance with the Declaration of Helsinki. All participants provided written informed consent and received a €50 food delivery service voucher as compensation. Measurements took place in June-September 2024.

### Stimuli and tasks

#### Artificial language stimuli

We adopted artificial language stimuli from Kuuluvainen et al. ^68^ to model early stages of language learning, where learners must segment continuous speech without clear word boundaries. SLL argues that the brain tracks transitional probabilities (TPs) between syllables to detect hidden regularities, enabling prediction of upcoming sensory stimuli. Each artificial language followed Finnish phonotactic and vowel harmony rules and comprised four trisyllabic words (TP = 1.0 within words; TP = 0.33 between words) presented without pauses or prosodic cues. Two distinct languages were used, each randomly assigned to one learning context (no-stress-induction vs. with-stress-induction). Each language stream lasted approximately two minutes, consisting of four 32-second exposure streams. Each word appeared 12 times per stream, totaling 48 repetitions across all four streams. For each language, four trisyllabic foils were created by recombining the same syllables into non-occurring sequences for later use in the 2AFC recognition test.

### Two-alternative forced choice (2AFC) and confidence rating task

We assessed auditory recognition of the artificial language using 2AFC task, a standard measure of SLL ^68,69^. Each trial presented a pair of trisyllabic items, one word from the exposure stream and one foil constructed from the same syllables in a novel order. Participants indicated which item “sounded more familiar” by clicking “Word 1” or “Word 2” on the screen. Trials were presented in random order, with word and foil positions counterbalanced across trials. After each decision, participants rated their confidence on a four-point scale (1 = guessing, 4 = remembering). Each trial began with a 500-ms fixation cross, followed by the first and second auditory items separated by a 1,500-ms inter-stimulus interval. Participants were instructed to respond quickly and intuitively and were informed that each item would be played only once. The SLL exposure streams and 2AFC tasks were implemented on the Gorilla Experiment Builder platform. The 2AFC task was administered twice: once for immediate testing following exposure, and again for delayed testing after the rest period (nap or wakeful rest).

### Virtual Reality – Trier Social Stress Test and Stroop Task

We induced acute psychosocial stress using the Finnish adaptation of the Virtual Reality Trier Social Stress Test (VR-TSST) ^43^, replicating the key elements of the original TSST which includes Job Interview and Subtraction tasks ^70^. We also supplemented the stress inductions with Virtual Reality Stroop Task ^44^. Participants completed the tasks while seated and wearing an HTC Vive Pro headset (HTC Corporation, Taoyuan, Taiwan).

The protocol included three components targeting complementary stress dimensions. The Job Interview elicited psychosocial-evaluative stress through simulated public speaking in front of a virtual panel. Participants delivered a five-minute speech explaining why they were ideal candidates for their “dream jobs” in front of three virtual judges. The Stroop task evoked cognitive–emotional interference by requiring color identification under incongruent word conditions. The Stroop task employed a standard color–word interference paradigm ^71^. The Subtraction task induced cognitive load and uncontrollability through time-pressured serial subtraction with performance feedback. Participants counted backward by thirteens from 1022, and in the second half by seventeens from 1033. Errors required restarting from the initial number to maintain performance pressure. The Job Interview and Stroop tasks were completed during the initial stress-induction phase, followed by a recovery period. The Subtraction task was administered after rest, near the end of the experiment.

### Recovery period

To approximate the natural dynamics of stress and recovery that occur in daily life, we extended the standard VR-TSST and VR-Stroop Test protocols so that they included longer recovery periods. Following the first stress session, participants completed a 15-minute Yoga Nidra relaxation while seated ^72^, followed by a 90-minute rest period involving either a nap or mind-wandering wakeful rest. Both groups underwent EEG recording during this period. Participants in the wakeful rest group engaged in quiet low-effort activities, coloring simple patterns and watching a silent nature documentary, and were instructed to let their minds wander freely without falling asleep. Participants in the nap group followed the standard daytime nap protocol under polysomnographic monitoring ^73^.

### Physiological measurement

#### Cardiac recording and Heart Rate Variability (HRV)

We recorded cardiac activity using a Polar H10 chest-strap sensor paired with a Polar Vantage V3 smart multi-sport watch (Polar Electro Oy, Kempele, Finland). The Polar H10 provides research-grade accuracy for R–R interval detection and has been validated against clinical electrocardiogram systems in both resting and dynamic conditions ^74^. Cardiac activity was recorded continuously while participants wore the sensor, with the watch acting as a receiver to log the data. Timestamps were recorded to mark the start and end of each experimental phase (baseline, stress induction tasks, post-stress task rebound, Yoga Nidra relaxation, nap/wakeful rest, language exposure, and the 2AFC task).

R–R interval data were processed in Kubios HRV Premium (version 3.5; Kubios Oy, Kuopio, Finland). The software applies automated artifact correction, trend removal, and beat-detection algorithms to clean the raw signal ^75^. We visually inspected all segments to confirm preprocessing accuracy.

In this study, the choice of R-R derived metrics was motivated by our aim to characterize individual differences in cardiac responses across an extended stress-recovery paradigm. Recent HRV publication guidelines recognize that there is no single gold-standard measure that should be adopted across all contexts, as the suitability of different metrics depends on the research question and experimental paradigm ^22^. Previous studies focusing on individual differences in autonomic cardiac regulation have used multiparametric composite indices to summarize physiological profiles, combining several features of the R-R signal ^50–53^. Following this approach, we adopted Kubios SNS and PNS indices as operational measures of cardiac mobilization (CM) and cardiac vagal recovery (CR), respectively. However, we refrain from making any claim that these indices are direct measures of SNS and PNS neural activity or can provide a branch-specific representation of autonomic cardiac control. We used these indices to provide a consistent multiparametric measure of changes in cardiac responses across the stress and recovery segments in the experiment.

The Kubios SNS index combines mean heart rate, Baevsky’s Stress Index (SI), and normalized Poincaré-plot SD2%. Increased heart rate is an established marker of cardiac mobilization, including due to acute psychosocial stress ^76^. SI has physiological content relevant to cardiac mobilization, as it quantifies the concentration and reduced dispersion of R–R intervals that typically accompany cardiovascular challenge ^77^. SD2 has been argued to reflect variation in cardiac mobilization, based on observations that it progressively changed following perturbations despite complete vagal blockade ^78^. Therefore, we concluded that the Kubios SNS index had relevant features for use as a proxy for our cardiac mobilization (CM) construct.

Meanwhile, the Kubios PNS index combines mean R-R interval, raw root mean square of successive differences (RMSSD), and normalized SD1%. The mean R-R interval captures cardiac slowing with longer intervals observed during recovery from acute psychological stress ^78^. RMSSD reflects predominantly vagally mediated beat-to-beat variation in heart rate intervals ^49^. Poincaré plot SD1, which represents the width of the Poincaré cloud, has been argued to indicate short-term HRV dominated by fast vagal influence ^78,79^. We thus concluded that the Kubios PNS index had relevant features to represent our cardiac vagal recovery (CR) construct.

The issue with Kubios indices is their proprietary weighting that limits transparency and reproducibility. We mitigated this issue by conducting a post-hoc linear reconstruction of the Kubios indices from their R-R derived components. We also conducted sensitivity analyses using alternative CM and CR operationalizations, following the triangulation recommendation from Quigley and colleagues ^22^. Full methods are provided in the Supplementary Material.

### Electrodermal activity (EDA)

We used EDA to record skin conductance level during the stress induction with galvanic skin sensors attached to the middle and index fingers of the non-dominant hand, connected to a QuickAmp amplifier (Brain Products GmbH, Gilching, Germany; sampling rate = 500 Hz). We analyzed the skin conductance (SC) signal as a whole, without decomposing it into tonic (SCL) and phasic (SCR) components. VR-TSST comprises extended, non-event-locked task segments for which block-wise averages of the SC signal provide a reliable index of sympathetic activation ^80^. Mean SC values were calculated for each task segment (e.g., Waiting Room, Job Interview, Stroop) and baseline periods, and stress reactivity was assessed by comparing mean SC during the stress segments with the baseline levels. We used EDA only to validate stress induction and did not include it in the composite score for cardiac mobilization.

### Procedure

The experiment lasted approximately six hours and was completed on a single laboratory visit (see Fig. 1 for graphical timeline). Before arrival, participants completed an electronic pre-study questionnaire assessing demographics, sleep quality, and general health. Upon arrival at the sleep laboratory (11:00 for wakeful rest group and 12:00 for nap group), they provided written informed consent and were fitted with physiological recording devices.

The 65 participants were semi-randomly assigned to either nap (*n* = 33) or wakeful rest (*n* = 32) and to one of two counterbalanced task-order groups, AB (*n* = 32) or BC (*n* = 33). The resulting allocation was nap–AB (*n* = 16), nap–BC (*n* = 17), wakeful-rest–AB (*n* = 16), and wakeful-rest–BC (*n* = 16). Following outlier exclusions, the final sample comprised 60 participants: nap–AB (*n* = 16), nap–BC (*n* = 14), wakeful-rest–AB (*n* = 15), and wakeful-rest–BC (*n* = 15).

The session began with baseline recordings of heart rate (HR) and EDA. Participants in the AB group (no-stress-induction first) completed the control SLL task (language exposure and immediate 2AFC), followed by acute stress inductions through the Job Interview component of VR-TSST and Stroop tasks. Approximately five minutes after the stress inductions, they then performed the with-stress-induction SLL task. Participants in the BC group (with-stress-induction first) completed the with-stress-induction SLL task after the stress inductions and their no-stress-induction SLL task after the subsequent recovery phase.

After the stress inductions, all participants entered the recovery period, beginning with a 15-minute guided Yoga Nidra relaxation, followed by either a 90-minute daytime nap or a mind-wandering wakeful rest. EEG and HR were continuously recorded throughout this period. Participants then had a snack break and shower before continuing the session. Following recovery, participants completed the delayed 2AFC test, after which acute stress was re-induced via the Subtraction task component of the VR-TSST. Subjective affect and stress evaluations were collected at six time points across the session, after each stress and recovery phase.

The session concluded around 17:00–18:00, after which all equipment was removed, and participants were debriefed. Counterbalancing ensured equal representation of nap versus wakeful rest groups and AB versus BC task orders. To maintain a focused scope, EEG and subjective affect data are reported in separate publications.

### Statistical analyses

We performed the analyses in Python (version 3.13; Python Software Foundation, 2024) and R (version 4.5.0; R Core Team, 2025).

### Group-level analysis

We assessed statistical learning accuracy by calculating the percentage of correctly identified words from all word–foil pairs, separately for each within-subject context (no-stress-induction and with-stress-induction) and phase (immediate and delayed 2AFC). Two-sided paired-samples *t*-tests in R were used to compare learning accuracy between the no-stress-induction and with-stress-induction learning contexts within each phase, and two-sided one-sample *t*-tests against chance level (50%) were used to determine whether accuracy exceeded random guessing.

### Composite cardiac response scores

We computed composite scores for cardiac mobilization (CM) and cardiac vagal recovery (CR) to capture individual differences in cardiac responses across stress and recovery contexts during the experiment. This approach was motivated by work showing that individual differences in cardiovascular reactivity can be more reliably characterized by integrating responses across multiple challenges than by relying on responses to a single task ^81,82^. Prior work has also shown that individual differences in autonomic cardiac control are highly correlated and stable across speech, arithmetic and reaction-time stress tasks ^18^. Accordingly, baseline-corrected CM and CR indices derived from both stress and recovery phases were entered into separate principal component analyses (PCA) to derive a composite score for each response dimension. Baseline 1, measured upon arrival at the laboratory, was used to correct all task measures except that from the Subtraction Task, which was corrected using Baseline 2, which was measured after the rest period (see Fig. 2). Analyses were performed in Python using *pandas*, *scikit-learn*, and *factor_analyzer*. All variables were z-standardized prior to PCA, sampling adequacy was confirmed using Bartlett’s Test of Sphericity and the Kaiser–Meyer–Olkin (KMO) measure, and component stability was evaluated via 1,000 bootstrap iterations. The first principal component (PC1) from each analysis was retained as a continuous composite score for use in subsequent mixed-effects models.

### Generalized linear mixed-effect modelling (GLMM)

To examine how cardiac responses predicted learning accuracy, we fitted a generalized linear mixed-effects model (GLMM) with a binomial distribution (Correct = 1, Incorrect = 0) and a logit link function using the lme4 package in R ^83^. Model comparisons were conducted using likelihood-ratio tests. This modelling choice followed recommendations for analyzing binary psycholinguistic data with repeated observations across participants and items ^84,85^, and is consistent with analytical approach in previous statistical learning studies ^68,86,87^.

All the categorical factors in the GLMM were effect-coded (e.g., −1 = no-stress-induction; +1 = with-stress-induction) so that coefficients reflected effects averaged across both categories ^88^. All continuous predictors were standardized (z-scored). Prior to model fitting, we confirmed that CM and CR scores were not strongly correlated (Pearson’s r = −0.053), indicating no problematic multicollinearity.

Model construction followed a theoretically motivated sequence for nested models. We initially allowed participants to differ in their accuracy and in the effects of Context (no-stress-induction vs. with-stress-induction) and Phase (immediate vs. delayed 2AFC testing), following recommendations to consider maximal random effects justified by the study design ^89^. We also included an item random intercept to account for differences in difficulty across the 2AFC word–foil test pairs ^84,86^. The initial random-effects structure was singular and required simplification. In the simplified model, the estimated participant-to-participant variation in the Phase effect was zero, so we omitted the participant random slope for Phase. The final random-effects structure used for the primary analysis was therefore (1 + Context || Participant) + (1 | Item), allowing participants to differ in accuracy and Context effects, and items to differ in difficulty. This model converged without a singular fit.

We entered fixed effects sequentially according to their theoretical relevance. We included covariates (Order Group, Rest Type, Age, and Gender) first, followed by task factors (Phase and Context) and their interaction. We then added the CM and CR composite scores and their interaction to capture coordination between the two cardiac response dimensions. The CM composite × CR composite interaction was specified a priori. This decision was based on Vagal Tank Theory ^40^, which proposes that cardiac reactivity during challenge and subsequent recovery may provide more information about adaptive regulation when considered jointly than when either response is considered alone. It was also motivated by Knight et al. ^90^ who applied Berntson’s two-dimensional Autonomic Space Model to examine how cardiac autonomic dimensions jointly predict cognitive functioning.

Next, momentary CM during language learning and its interaction with CM composite were introduced to test whether momentary cardiac mobilization during speech exposure interacted with the individual’s characteristic cardiac mobilization across all stress induction tasks. The inclusion of both momentary CM and characteristic CM composite predictors was theoretically motivated by integrative frameworks linking a relatively stable physiological differences with dynamic regulatory responses ^91^.

To examine whether the associations between the CM and CR composites and learning accuracy differed between immediate and delayed testing, we included the Phase × CM composite × CR composite interaction and its lower-order terms. We also tested higher-order interactions with Context, but these did not improve model fit and were not retained. Furthermore, we tested the Phase × Order Group interaction because the counterbalanced order groups (AB and BC) differed in the average elapsed time between immediate and delayed testing. We also tested the Phase × Rest Type to see whether the change in learning performance from immediate to delayed testing differed between nap and wakeful rest. These interactions likewise did not improve model fit and were not included in the final model. To avoid overfitting, only interactions that were theoretically relevant to the research questions were evaluated.

The final model structure was *Accuracy ∼ Phase + Context + Covariates + (CM composite × CR composite) + (CM composite × momentary CM) + (Phase × CM composite × CR composite) + (1 + Context || Participant) + (1 | Item)*.

Model assumptions were evaluated using residual diagnostics (Q–Q plots, residuals-versus-fitted inspection, binned residual tests with DHARMa) and checks for overdispersion. Predicted marginal effects were visualized with ggeffects and ggplot2, with 95% confidence intervals and standardized (±1 SD) values for interpretability.

In post-hoc sensitivity analyses, we also tested baseline CM and baseline CR measurement as additional covariate.

### Cardiac response profile quadrant

To provide intuitive visualization of the combined effects of CM and CR on learning and to create categorical profiles of cardiac response congruence, we divided participants’ z-standardized CM and CR composite scores at their respective sample means.

## Author contributions: CRediT

**Aliva Sholihat:** Conceptualization, Data curation, Formal analysis, Investigation, Methodology, Project administration, Software, Visualization, Writing – original draft. **Risto Halonen**: Conceptualization, Data curation, Investigation, Methodology, Project administration, Resources, Software, Supervision, Validation, Writing – review and editing. **Riikka Möttönen**: Conceptualization, Funding acquisition, Methodology, Resources, Supervision, Validation, Writing – review and editing. **Anu-Katriina Pesonen**: Conceptualization, Funding acquisition, Methodology, Resources, Supervision, Validation, Writing – review and editing.

## Acknowledgements

We thank Kaisu Martinmäki from Polar Electro Oy, for collaboration in the data collection; research assistants Elisa Kotala and Senni Sipponen for practical management of the experiment; Soila Kuuluvainen, Saara Kaskivuo and Martti Vainio for the design of the artificial language stimuli.

This work was supported by Research Council of Finland grants no. 1356020 (to A-K.P.) and 3142979 (to R.M.). R.M. was also supported by the Strategic Research Council (373227). A.S. was supported by a fellowship from the Doctoral School of the University of Helsinki. The funding body was not involved in the study design, data collection, data analysis, publication decision, or manuscript preparation. The views, findings, conclusions presented here are solely those of the authors and do not necessarily represent those of the funders.

## Data availability

The anonymized datasets used in the manuscript and the analysis code are available on Open Science Framework platform. The link will be provided in the published version of the manuscript.

## Declaration of generative AI and AI-assisted technologies in the manuscript preparation process

During the preparation of this work the authors used AI-assisted technologies (ChatGPT and Claude) for language support, including grammar, phrasing, and refinement. After using these tools, the authors reviewed and edited the content as needed and took full responsibility for the content of the published article.

