## Supplementary_material for "Language learning in a stressful world: individual differences in cardiac responses to stress and recovery predict statistical learning"

Sholihat et al., 2026

**This document includes:**

**Main analysis**

Table S1

**Post-hoc analyses**

Fig. S1 to S2

Method Fig. S1

Tables S2 to S6

Method Table S2 to S3

**Main analysis**

Table S1.

GLMM results with Kubios-derived cardiac mobilization (CM) and cardiac vagal recovery (CR)

| **Predictor / contrast** | **Odds ratio¹** | **95% CI (Low)** | **95% CI (High)** | **p-value** |
| --- | --- | --- | --- | --- |
| **Intercept** | **1.37** | **1.09** | **1.73** | **0.007** |
| **Phase (delayed vs. immediate)** | **0.74** | **0.65** | **0.85** | **< .001** |
| Learning context (with-stress-induction vs. no-stress-induction) | 1.15 | 0.93 | 1.43 | 0.206 |
| Order group (AB vs. BC) | 0.90 | 0.75 | 1.08 | 0.271 |
| Rest type (wakeful rest vs. nap) | 0.99 | 0.84 | 1.16 | 0.870 |
| Age (standardized) | 1.06 | 0.97 | 1.15 | 0.184 |
| Gender (female vs. male) | 1.00 | 0.83 | 1.19 | 0.973 |
| CM composite | 1.03 | 0.95 | 1.13 | 0.468 |
| CR composite | 1.07 | 0.98 | 1.16 | 0.154 |
| Momentary CM | 0.98 | 0.88 | 1.09 | 0.692 |
| **CM composite × CR composite** | **1.15** | **1.05** | **1.25** | **0.002** |
| **CM composite × Momentary CM** | **1.16** | **1.06** | **1.27** | **< .001** |
| **Phase × CM composite (delayed vs. immediate)** | **0.83** | **0.72** | **0.95** | **0.008** |
| **Phase × CR composite (delayed vs. immediate)** | **1.17** | **1.02** | **1.34** | **0.025** |
| Phase × CM composite × CR composite (delayed vs. immediate) | 1.09 | 0.95 | 1.24 | 0.211 |

N = 60 participants; 3840 responses; 32 items. AIC = 4944.45; BIC = 5057.01.

Note. ¹ Binary-category contrasts and phase interaction contrasts use exp(2β), reflecting the original −1/+1 coding; other rows use exp(β). Interactions are ratios of odds ratios, and the intercept is odds. Confidence intervals are 95% Wald intervals. Bold rows indicate p < .05.

**Post-hoc sensitivity analyses**

Fig. S1.

Reconstruction of Kubios SNS and PNS indices with participant-held-out linear regression model. Each data point represents one eligible segment of SNS and PNS indices. Dashed lines indicate near perfect agreement of the reconstructed indices and the original indices. Equation above the panels are the final full-sample regression equations. Plotted predictions and performance charts come from five-fold participant held-out validation. Metrics used are: HR (bpm), raw RMSSD (ms), SD1% and SD2% (in percentage share), SI uses the Kubios-exported scale.

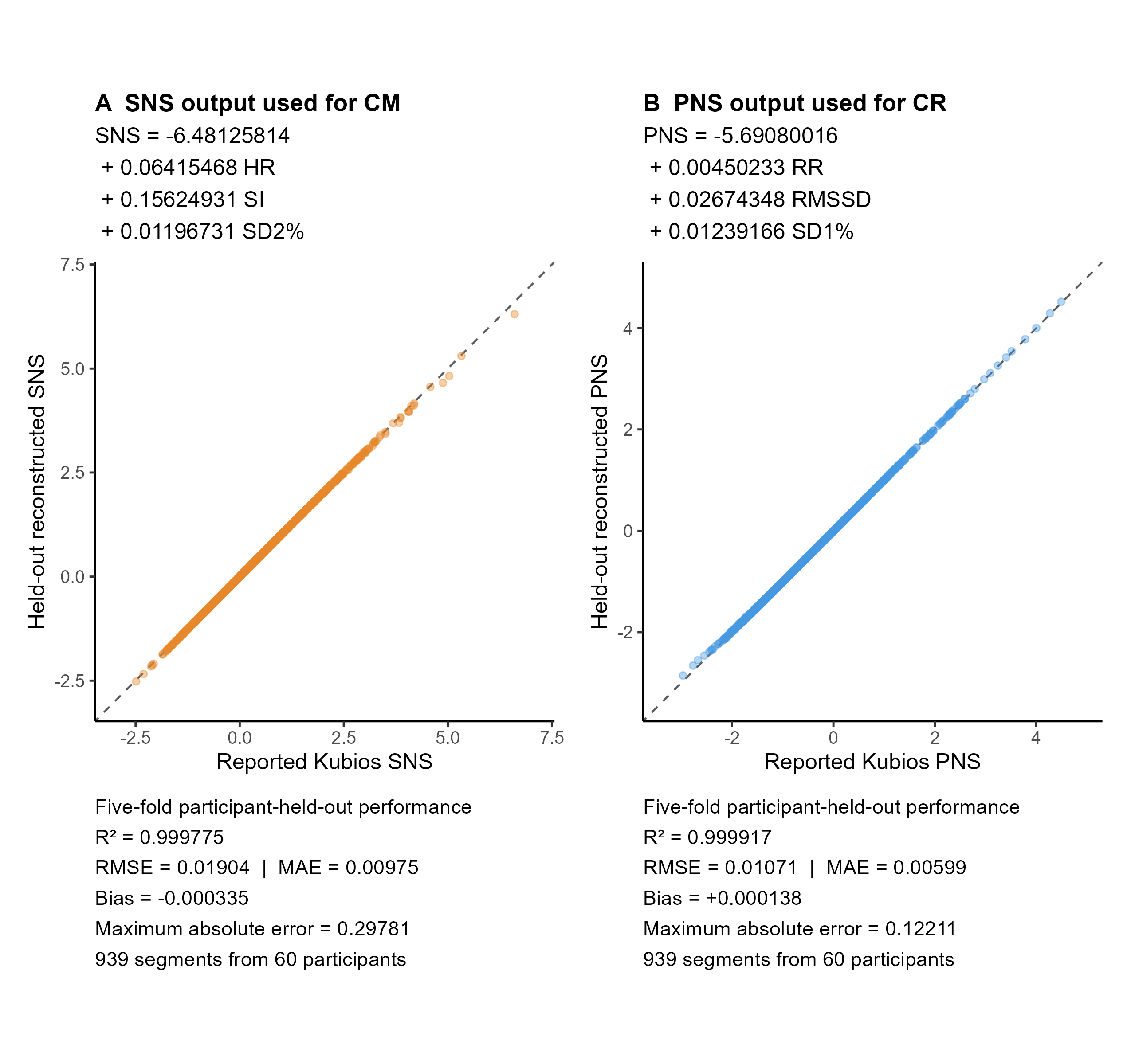

Method Fig. S1

**Transparent linear reconstruction of Kubios indices**

In this linear regression model, we examined whether we could reconstruct the Kubios indices using their publicly documented components. The aim was to reproduce the numerical values and conclude from the model parameters whether their proprietary weighting was essential to our findings.

The analysis used data from the clean 60 participants after outlier exclusion. We recorded 960 segments, with 939 containing the necessary numerical values of the R-R component. Fifty-six participants contributed 16 segments each, while the remaining four participants contributed 15, 14, 12, and 2 segments respectively.

For the reconstructed SNS index, the equation included HR (bpm), stress index (SI), and SD2%. For the PNS index, the equation included mean RR (ms), raw RMSSD (ms), and SD1%. These components were based on publicly documented Kubios description of their indices [1].

We calculated SD1% as 100 × SD1/(SD1 + SD2). Meanwhile, SD2% was calculated as 100 × SD2/(SD1 + SD2). SI used the direct numerical values as exported from Kubios without applying any additional log or square-root transformation, following the Kubios description of Beavsky’s SI [2].

We fitted a separate weighted linear regression for each SNS and PNS indices, with the three components as predictors. We included weight for each segment by 1 divided by the number of eligible segments from that participant to ensure that participants with more segments did not contribute disproportionately to the overall weight of the regression. To estimate the uncertainty around the regression coefficients, we used 2,000 bootstrap samples. We then evaluated the equations using participant-level five-fold cross-validation. In each round, we fitted the equations on 48 participants and predicted Kubios values for the remaining 12 and repeated the process five times. We then combined all test predictions and assessed agreement with Kubios values using R², RMSE (root mean square error), MAE (mean absolute error), signed bias (predicted minus reported values), and maximum absolute error.

Finally, we applied the fitted regression equations to sample data from the publicly available Kubios HRV user guide [3] as a small external check. The predicted SNS and PNS indices closely matched the reported values, with all absolute differences below 0.02 index units. Nevertheless, further validation using larger independent datasets is needed to assess whether these transparent equations can consistently reproduce Kubios values across different samples and recording conditions.

All reconstruction analyses were conducted in R version 4.5.0. Weighted linear regressions were fitted using the lm() function in the stats package, with participant-level bootstrap refitting performed using lm.wfit().

Fig. S2.

Physiological dynamics during the experiment across different operationalizations of CM and CR.

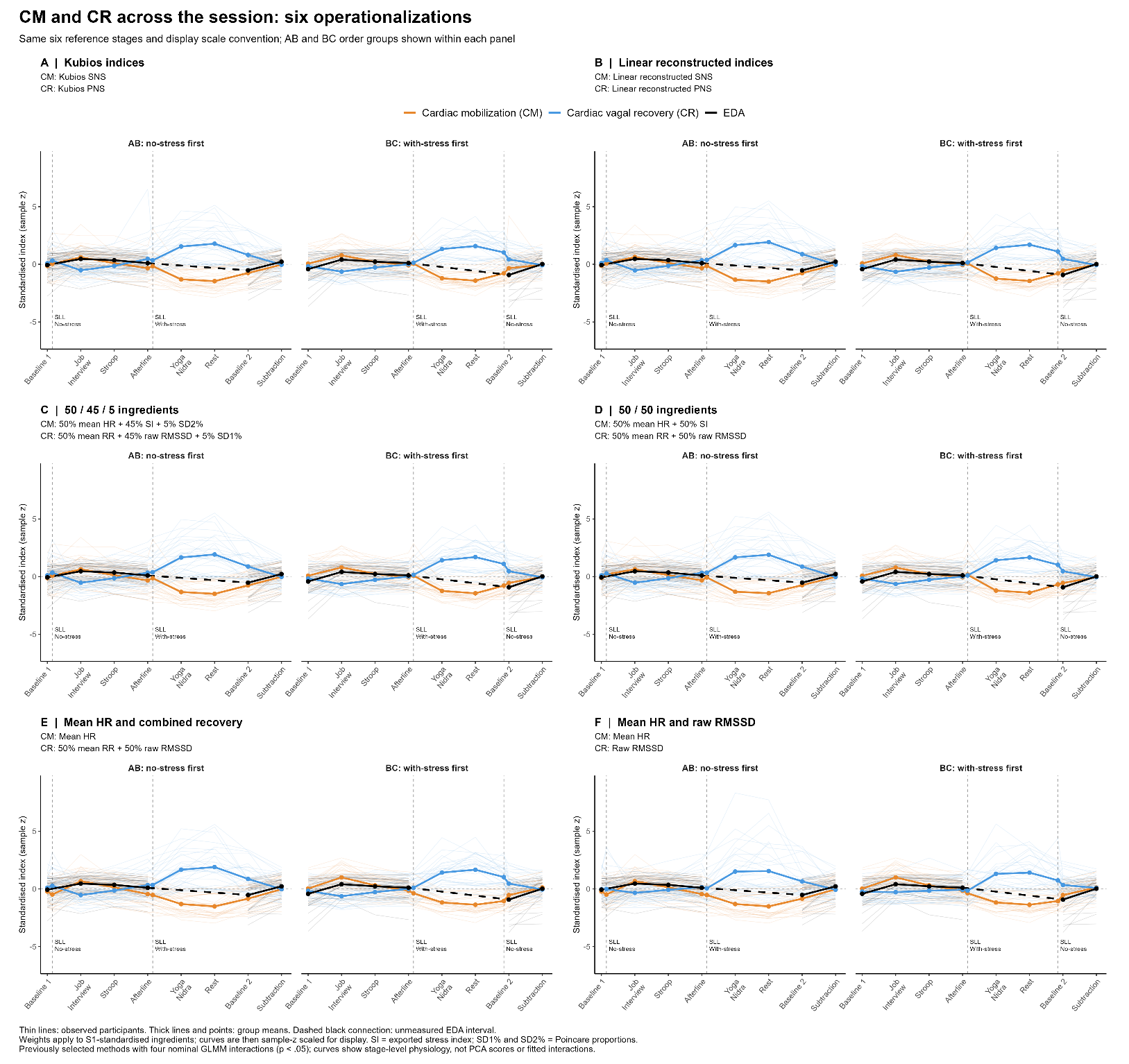

Table S2.

Model fitting with different operationalization of cardiac mobilization (CM) and cardiac vagal recovery (CR), summary of p-values

| **CM x CR operationalization** | **CM × CR** | **CM × Momentary CM** | **CM × Phase** | **CR × Phase** | **AIC** | **BIC** |
| --- | --- | --- | --- | --- | --- | --- |
| **(Kubios SNS) x (Kubios PNS)** | **.0021 **** | **<.001 ***** | **.0082 **** | **.0249 *** | **4944.45** | **5057.01** |
| **Linear regression reconstructed Kubios from transparent components** | **.0219 *** | **.0096 **** | **.0109 *** | **.0206 *** | **4947.19** | **5059.75** |
| **(50% mean HR + 45% SI + 5% SD1) x (50% mean RR + 45% raw RMSSD + 5% SD2)** | **.0233 *** | **.0093 **** | **.0110 *** | **.0194 *** | **4947.14** | **5059.70** |
| (50% mean HR + 45% SI + 5% SD1) x (50% mean RR + 45% lnRMSSD + 5% SD2) | .0492 * | .0088 ** | .0157 * | .0716 | 4950.59 | 5063.15 |
| **(50% mean HR + 50% SI) x (50% mean RR + 50% raw RMSSD)** | **.0471 *** | **.0087 **** | **.0137 *** | **.0118 *** | **4947.30** | **5059.86** |
| (50% mean HR + 50% SI) x (50% mean RR + 50% lnRMSSD) | .0935 | .0080 ** | .0220 * | .0439 * | 4951.20 | 5063.76 |
| **(Mean HR) x (50% mean RR + 50% raw RMSSD)** | **.0027 **** | **.0107 *** | **.0440 *** | **.0192 *** | **4948.96** | **5061.51** |
| (SI) x (50% RR + 50% raw RMSSD) | .2245 | .0121 * | .0304 * | .0136 * | 4949.00 | 5061.56 |
| (Mean HR) x (mean RR) | .00085 *** | .0109 * | .0336 * | .3834 | 4952.80 | 5065.36 |
| **(Mean HR) x (raw RMSSD)** | **.0214 *** | **.0391 *** | **.0409 *** | **.0048 **** | **4949.55** | **5062.11** |
| (Mean HR) x (lnRMSSD) | .2042 | .0685 | .1003 | .0108 * | 4954.37 | 5066.93 |
| (lnSI) x (lnRMSSD) | .7838 | .1085 | .0894 | .0030 ** | 4954.26 | 5066.82 |
| (Kubios SNS) x (lnRMSSD) | .3128 | .0188 * | .0312 * | .0096 ** | 4950.95 | 5063.51 |
| (lnRMSSD withdrawal) x (lnRMSSD recovery) | .7091 | .5615 | .1261 | .0062 ** | 4955.78 | 5068.33 |
| (mean HR) x (lnRMSSD) | .2120 | .0704 | .1025 | .0082 ** | 4954.33 | 5066.89 |
| (Kubios SNS) x (raw RMSSD) | .0538 | .0109 * | .0085 ** | .0033 ** | 4945.83 | 5058.39 |

Method Table S2

**Sensitivity to alternative operationalizations of cardiac mobilization and cardiac vagal recovery**

We investigated whether associations between cardiac mobilization (CM), cardiac vagal recovery (CR), and learning accuracy depended on their physiological operationalization. The main aim was to determine if the study’s findings depended on the proprietary Kubios-derived indices which might limit transparency and reproducibility.

Different alternatives tested included: Kubios SNS/PNS indices, linear reconstructions from their documented components, weighted combinations of their components, and individual metric of mobilization and recovery using either their raw format or the log-naturalized format. Each alternative operationalization was first fitted to the PCA step to derive the CM and CR composite indices. The CM and CR composite indices were then fitted to the final model.

Each operationalization was evaluated using the same final structure of the binomial generalized linear mixed model (GLMM), retaining all 3,840 responses from 60 participants and 32 items:

Accuracy ~ Phase + Context + Covariates + (CM_composite * CR_composite) + (CM_composite * momentary_CM) + (Phase * CM_composite * CR_composite) + (1 + Context || Participant) + (1 | Item)

Accuracy was coded as correct or incorrect. Phase distinguished learning (−1) from recall (+1), Context denoted the experimental condition, and Covariates comprised condition order, rest type, standardized age, and gender.

Models were fitted by maximum likelihood using the Laplace approximation in R’s lme4 package.

Table S3.

Model fitting with different operationalizations of cardiac mobilization (CM) and cardiac vagal recovery (CR), summary of p-values.

Includes the six operationalizations that yielded all four significant interactions in models without baseline covariates (see Table 2 in bold). Results below include initial-baseline covariates.

| **CM × CR operationalization** | **CM × CR** | **CM × Momentary CM** | **CM × Phase** | **CR × Phase** | **Baseline CM** | **Baseline CR** | **AIC** | **BIC** |
| --- | --- | --- | --- | --- | --- | --- | --- | --- |
| **Kubios SNS × Kubios PNS** | **0.0103 *** | **0.0031 **** | **0.0083 **** | **0.0250 *** | **0.3135** | **0.5557** | **4945.84** | **5070.90** |
| Linear regression reconstructed Kubios from its components | 0.0625 | 0.0242 * | 0.0111 * | 0.0206 * | 0.1979 | 0.4007 | 4947.73 | 5072.79 |
| (50% mean HR + 45% SI + 5% SD2%) × (50% mean RR + 45% raw RMSSD + 5% SD1%) | 0.0636 | 0.0227 * | 0.0112 * | 0.0194 * | 0.2435 | 0.4667 | 4947.87 | 5072.93 |
| (50% mean HR + 50% SI) × (50% mean RR + 50% raw RMSSD) | 0.1064 | 0.0209 * | 0.0139 * | 0.0119 * | 0.2383 | 0.5040 | 4947.69 | 5072.76 |
| **Mean HR × (50% mean RR + 50% raw RMSSD)** | **0.0045 **** | **0.0306 *** | **0.0444 *** | **0.0192 *** | **0.0832** | **0.3062** | **4949.07** | **5074.13** |
| Mean HR × raw RMSSD | 0.0267 * | 0.0812 | 0.0411 * | 0.0048 ** | 0.0411 * | 0.4013 | 4949.47 | 5074.54 |

Only the original Kubios and mean HR × equal-weight mean RR/raw RMSSD models retained all four significant interactions after baseline adjustment.

Method Table S3:

**Sensitivity of CM and CR interactions to adjustment for initial resting cardiac levels**

To investigate whether the observed interactions remained after accounting for initial resting cardiac levels, we chose the six operationalizations in Table S2 which all the four interactions were significant.

Each GLMM was refitted with two additional covariates representing initial resting baseline of CM and CR.

The adjusted model specification in R notation was:

Accuracy ~ Phase + Context + Covariates + Baseline_CM + Baseline_CR + (CM_composite * CR_composite) + (CM_composite * momentary_CM) + (Phase * CM_composite * CR_composite) + (1 + Context || Participant) + (1 | Item)

Models were fitted by maximum likelihood using the Laplace approximation in R’s lme4 package.

Table S4.

GLMM results with Kubios-derived cardiac mobilization (CM) and cardiac vagal recovery (CR)

*With initial-baseline covariates*

| **Predictor / contrast** | **Odds ratio¹** | **95% CI (Low)** | **95% CI (High)** | **p-value** |
| --- | --- | --- | --- | --- |
| **Intercept** | **1.38** | **1.10** | **1.74** | **0.006** |
| **Phase (delayed vs. immediate)** | **0.74** | **0.65** | **0.85** | **< .001** |
| Learning context (with-stress-induction vs. no-stress-induction) | 1.15 | 0.93 | 1.43 | 0.200 |
| Order group (AB vs. BC) | 0.90 | 0.75 | 1.08 | 0.248 |
| Rest type (wakeful rest vs. nap) | 0.94 | 0.79 | 1.12 | 0.504 |
| Age (standardized) | 1.06 | 0.97 | 1.15 | 0.186 |
| Gender (female vs. male) | 0.98 | 0.83 | 1.17 | 0.862 |
| CM composite | 1.05 | 0.96 | 1.15 | 0.270 |
| CR composite | 1.08 | 0.99 | 1.18 | 0.073 |
| Momentary CM | 0.98 | 0.88 | 1.10 | 0.773 |
| **CM composite × CR composite** | **1.12** | **1.03** | **1.23** | **0.010** |
| **CM composite × Momentary CM** | **1.15** | **1.05** | **1.25** | **0.003** |
| **Phase × CM composite (delayed vs. immediate)** | **0.83** | **0.72** | **0.95** | **0.008** |
| **Phase × CR composite (delayed vs. immediate)** | **1.17** | **1.02** | **1.34** | **0.025** |
| Phase × CM composite × CR composite (delayed vs. immediate) | 1.09 | 0.95 | 1.24 | 0.211 |
| Baseline CM (standardized) | 1.17 | 0.86 | 1.57 | 0.313 |
| Baseline CR (standardized) | 1.09 | 0.82 | 1.46 | 0.556 |

N = 60 participants; 3840 responses; 32 items. AIC = 4945.84; BIC = 5070.90.

Note. ¹ Binary-category contrasts and phase interaction contrasts use exp(2β), reflecting the original −1/+1 coding; other rows use exp(β). Interactions are ratios of odds ratios, and the intercept is odds. Confidence intervals are 95% Wald intervals. Bold rows indicate p < .05.

Table S5.

GLMM results with mean HR as proxy for cardiac mobilization (CM) and equally weighted mean RR and raw RMSSD as proxy for cardiac vagal recovery (CR)

| **Predictor / contrast** | **Odds ratio¹** | **95% CI (Low)** | **95% CI (High)** | **p-value** |
| --- | --- | --- | --- | --- |
| **Intercept** | **1.38** | **1.10** | **1.75** | **0.006** |
| **Phase (delayed vs. immediate)** | **0.74** | **0.65** | **0.85** | **< .001** |
| Learning context (with-stress-induction vs. no-stress-induction) | 1.14 | 0.93 | 1.41 | 0.217 |
| Order group (AB vs. BC) | 0.95 | 0.78 | 1.16 | 0.621 |
| Rest type (wakeful rest vs. nap) | 0.93 | 0.79 | 1.11 | 0.432 |
| Age (standardized) | 1.08 | 0.99 | 1.18 | 0.098 |
| Gender (female vs. male) | 1.00 | 0.83 | 1.21 | 0.966 |
| CM composite | 0.97 | 0.89 | 1.06 | 0.479 |
| CR composite | 1.06 | 0.97 | 1.17 | 0.196 |
| Momentary CM | 0.92 | 0.82 | 1.05 | 0.207 |
| **CM composite × CR composite** | **1.16** | **1.05** | **1.27** | **0.003** |
| **CM composite × Momentary CM** | **1.14** | **1.03** | **1.26** | **0.011** |
| **Phase × CM composite (delayed vs. immediate)** | **0.87** | **0.75** | **1.00** | **0.044** |
| **Phase × CR composite (delayed vs. immediate)** | **1.18** | **1.03** | **1.35** | **0.019** |
| Phase × CM composite × CR composite (delayed vs. immediate) | 1.06 | 0.93 | 1.21 | 0.403 |

N = 60 participants; 3840 responses; 32 items. AIC = 4948.96; BIC = 5061.51.

Note. ¹ Binary-category contrasts and phase interaction contrasts use exp(2β), reflecting the original −1/+1 coding; other rows use exp(β). Interactions are ratios of odds ratios, and the intercept is odds. Confidence intervals are 95% Wald intervals. Bold rows indicate p < .05.

Table S6.

GLMM results with mean HR as proxy for cardiac mobilization (CM) and equally weighted mean RR and raw RMSSD as proxy for cardiac vagal recovery (CR)

*With initial-baseline covariates*

| **Predictor / contrast** | **Odds ratio¹** | **95% CI (Low)** | **95% CI (High)** | **p-value** |
| --- | --- | --- | --- | --- |
| **Intercept** | **1.39** | **1.10** | **1.75** | **0.006** |
| **Phase (delayed vs. immediate)** | **0.74** | **0.65** | **0.85** | **< .001** |
| Learning context (with-stress-induction vs. no-stress-induction) | 1.15 | 0.93 | 1.42 | 0.202 |
| Order group (AB vs. BC) | 0.94 | 0.77 | 1.14 | 0.521 |
| Rest type (wakeful rest vs. nap) | 0.91 | 0.77 | 1.08 | 0.267 |
| **Age (standardized)** | **1.10** | **1.00** | **1.20** | **0.040** |
| Gender (female vs. male) | 1.02 | 0.85 | 1.22 | 0.816 |
| CM composite | 0.97 | 0.89 | 1.06 | 0.533 |
| CR composite | 1.07 | 0.97 | 1.19 | 0.151 |
| Momentary CM | 0.95 | 0.84 | 1.08 | 0.454 |
| **CM composite × CR composite** | **1.15** | **1.04** | **1.26** | **0.004** |
| **CM composite × Momentary CM** | **1.12** | **1.01** | **1.24** | **0.031** |
| **Phase × CM composite (delayed vs. immediate)** | **0.87** | **0.75** | **1.00** | **0.044** |
| **Phase × CR composite (delayed vs. immediate)** | **1.18** | **1.03** | **1.35** | **0.019** |
| Phase × CM composite × CR composite (delayed vs. immediate) | 1.06 | 0.93 | 1.21 | 0.403 |
| Baseline mean HR (standardized) | 1.18 | 0.98 | 1.42 | 0.083 |
| Baseline RR–RMSSD index (standardized) | 1.11 | 0.91 | 1.34 | 0.306 |

N = 60 participants; 3840 responses; 32 items. AIC = 4949.07; BIC = 5074.13.

Note. ¹ Binary-category contrasts and phase interaction contrasts use exp(2β), reflecting the original −1/+1 coding; other rows use exp(β). Interactions are ratios of odds ratios, and the intercept is odds. Confidence intervals are 95% Wald intervals. Bold rows indicate p < .05.
